# Unpredictable repeatability in molecular evolution

**DOI:** 10.1101/2022.05.24.493242

**Authors:** Suman G Das, Joachim Krug

## Abstract

The extent of parallel evolution at the genotypic level is quantitatively linked to the distribution of beneficial fitness effects (DBFE) of mutations. The standard view, based on light-tailed distributions (i.e. distributions with finite moments), is that the probability of parallel evolution in duplicate populations is inversely proportional to the number of available mutations, and moreover that the DBFE is sufficient to determine the probability when the number of available mutations is large. Here we show that when the DBFE is heavy-tailed, as found in several recent experiments, these expectations are defied. The probability of parallel evolution decays anomalously slowly in the number of mutations or even becomes independent of it, implying higher repeatability of evolution. At the same time, the probability of parallel evolution is *non-self-averaging*, that is, it does not converge to its mean value even when a large number of mutations are involved. This behavior arises because the evolutionary process is dominated by only a few mutations of high weight. Consequently, the probability varies widely across systems with the same DBFE. Contrary to the standard view, the DBFE is no longer sufficient to determine the extent of parallel evolution, making it much less predictable. We illustrate these ideas theoretically and through analysis of empirical data on antibiotic resistance evolution.

## I. INTRODUCTION

The repeatability and predictability of evolution are important questions in the field of evolutionary biology. In 1990, Stephen Jay Gould famously mused about replaying life’s tape [1]. In subsequent years, the topic of parallel evolution has become a major subject of empirical research [2–4], and theoretical questions concerning the probability of parallel evolution within the mathematical theory of population genetics have also attracted substantial attention [5–7]. Here the questions are focused mostly on changes at the level of genetic sequences. According to a common definition [6, 7], parallel evolution is said to occur when the exact same mutation is substituted in replicate populations. It is in this strong sense that we shall use the term *parallel evolution* here.

The computation of the probability of parallel evolution is often set in a simplified scenario [5–7] where an asexual population evolves by strong selection and weak mutation (SSWM); the evolutionary process starts with a homogeneous population, and any one of *n* available beneficial mutations, with selection coefficients *s_i_, i* = 1, 2,…, *n* can be the first to fix in the population. For small values of the *s_i_*, the probability that the *i*-th mutation will be the first to fix is *W_i_* = *s_i_*/∑*_j_ s_j_*) (see SI for more details). Therefore the probability that *k* replicate populations will all fix the same mutation is

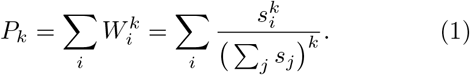

The mean probability of parallel evolution is 〈*P_k_*〉 where 〈·〉 denotes the average with respect to the DBFE. The set of selection coefficients varies across systems, but is expected to show statistical regularities [8]. It has been hypothesized [5–7] that, since viable organisms are already relatively well-adapted, mutants of higher fitness must be chosen from the tails of fitness distributions, and therefore the DBFE *P_s_* (*s*) must have the form of a limiting distribution in extreme-value theory (EVT). The EVT hypothesis predicts that the DBFE belongs to one of three classes of distributions: the Weibull and Gumbel classes contain distributions with finite moments, whereas the Fréchet class contains distributions with power law tails (and therefore diverging moments). In the last case, the asymptotic form of the DBFE is

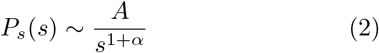

with a scale parameter *A* > 0 and the tail exponent *α* > 0. The cases of the Gumbel and Weibull extreme-value distributions have been explored in some detail for *k* = 2 [5–7]. The Frechet EVT class had been conjectured to be relatively unimportant biologically [7], but several subsequent studies [9–12] have uncovered signatures of heavy-tailed distributions of fitness effects (see SI for further details on tails of empirical DBFEs). In the realistic case of a large number of available beneficial mutations *n*, the statistics of *P_k_* for heavy-tailed distributions of the form (2) is markedly different from that of light-tailed distributions, as will be shown below. Note that in obtaining (1) we have taken the mutation rate *μ* to be uniform across mutations. When mutation bias is involved, one needs to make the replacement *μ* → *μs* in (1), and the analysis below will be applicable to heavy-tailed distributions of *μs* [2, 3].

## II. THEORY

The number *n* of beneficial mutations that can occur in a population varies widely across organisms and environments, but it is likely to be large. For bacterial populations, one may conservatively estimate that there are several thousand beneficial mutations (see SI). We therefore focus on *P_k_* in the large-*n* regime. The simplest computation is in the limiting case of neutral variation where all the selection coefficients are identical. Since all mutations are equally likely to be the first to fix, *P_k_* = 1/*n*^*k*-1^ (which is exact for all *n*). Specifically, the probability of parallel evolution in two replicates is *P*_2_ = 1/*n*. Using this observation, one can define, for any system, the quantity 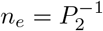 as the *effective number of mutations* that contribute to parallel evolution. It can be interpreted as the number of mutations in a different system which has the same *P*_2_ but where all the mutations are equally likely to fix in the population. Therefore *n_e_* is a measure of the number of mutations that dominate the dynamics of fixation. It is similar to the notion of the effective number of reproducing lineages studied in [13] in the context of family size distributions.

The most commonly studied class of DBFEs is where all the moments are finite. In this case, the numerator and denominator in each term in (1) become uncorrelated as *n* → ∞ and the distribution of 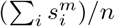 becomes sharply centred around the moment 〈*s^m^*〉 of *P_s_* (·). Therefore, for large *n*, we have

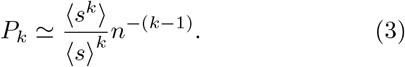

The solid brown line in Fig 1(a) shows this behavior of *P_k_* for the exponential distribution. Notice that so far we have omitted the angular brackets around *P_k_*, since it converges to the mean value in the limit of large *n*, as shown by the highly localized distribution of *P_k_* in Fig 1(b) (red dashed curve). Such quantities are described as *self-averaging*. For the particular case *k* = 2, we have 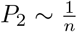, which is the characteristic decay in self-averaging systems. It was shown in [6] that for an exponential distribution, 〈*P_2_*〉 = 2/(*n*+1) (see SI for a general expression for 〈*P_k_*〉). Our focus here, however, is on heavy-tailed distributions with tails of the form (2). When *k* < *α*, (3) continues to hold. Particularly for *α* > 2, *P*_2_ still decays as ~ 1/*n*, as shown in Fig 1(a). However, when *k* > *α*, the moment 〈*s^k^*〉 diverges and (3) no longer holds. We can now break the analysis down into two cases.

**FIG. 1:**
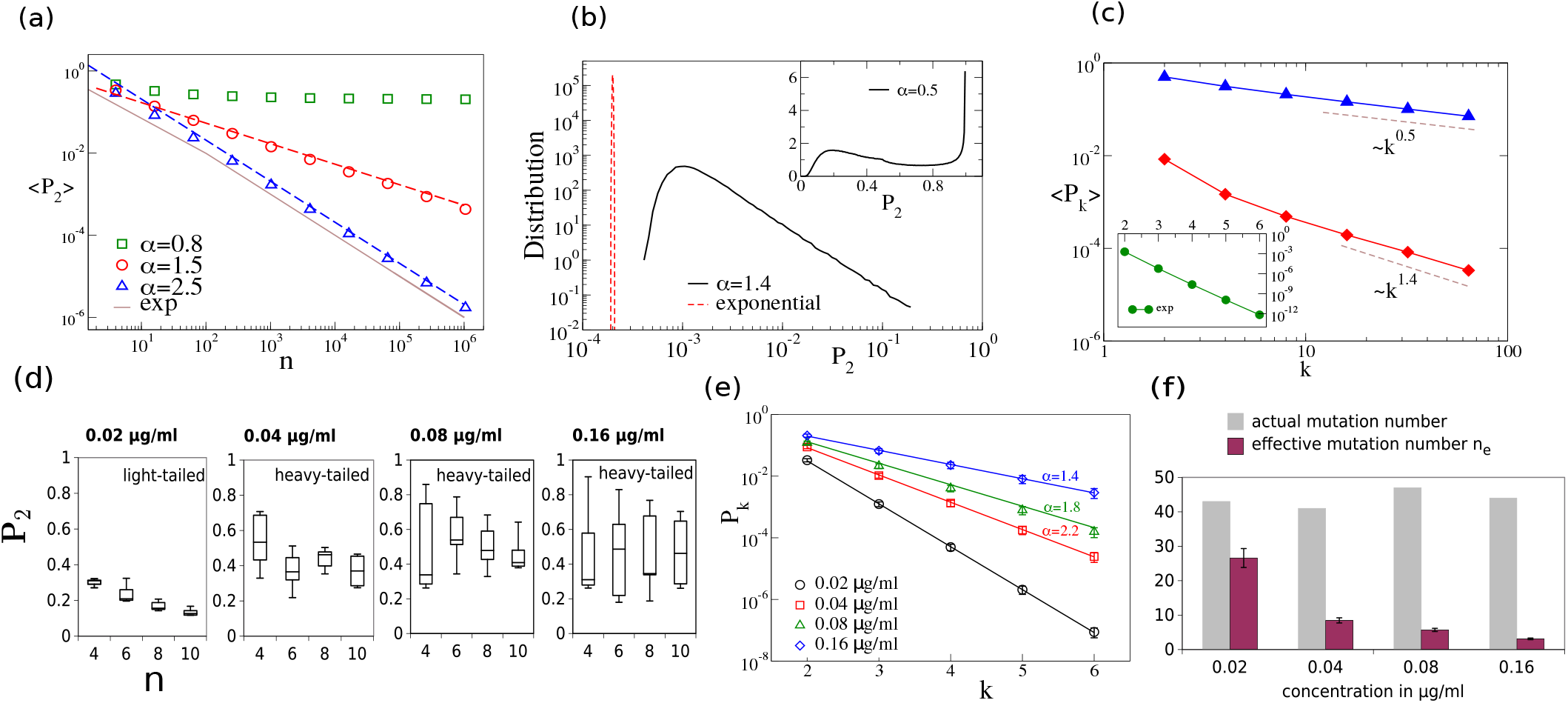
(a) Plot of (*P*_2_) for three different values of *α*. The symbols are numerically generated data with averages over 10^5^ realizations, and the dashed lines are the theoretical predictions from (4) and (6). The solid brown curve is the exact result for the exponential distribution. The curve for *α* = 2.5 has the same asymptotic behavior *P*_2_ ~ *n*^-1^ as that of the exponential since *α* > *k* = 2. (b) The black curve is the numerically sampled distributions of *P*_2_ for *α* = 1.4 and the inset shows the same for *α* = 0.5; we used 10^6^ realizations and *n* = 10^4^ mutations. Tire dashed red curve is the distribution of *P*_2_ for an exponential distribution of selection coefficients and *n* = 10^4^. (c) (*P_k_*) as a function of *k*, from (4) and (6). We used *n* = 10^4^ and 10^5^ realizations. The inset shows the rapid decay of (*P_k_*) for an exponential distribution and *n* = 10^3^, plotted using the exact result in SI. (d) This and the following figures analyze data from tlie study reported in [9] which determined tire selection coefficients for several mutations in TEM-1 *β*-lactamase conferring resistance to the antibiotic cefotaxime. Here we show the distribution of *P*_2_. The data set at each cefotaxime concentration was randomly split into subsets of size n. The box plots show median, quartiles, and extreme values. (e) The *P_k_* were obtained from the entire data set at each concentration and (5) were used to infer *α*. (f) The effective mutation number *n_e_* = 1/*P*_2_ has been computed and compared with the actual number of mutations in the available dataset at each concentration.

### Case I

The moderately heavy-tailed case occurs when *α* > 1; in this case 〈*s*〉 is finite, but higher moments corresponding to *k* > *α* > 1 diverge. For *k* > *α*, the asymptotic behavior of (*P_k_*) is

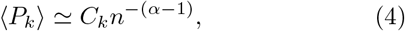

where the constant *C_k_* = *A*Γ(*k – *α**)Γ(*α*)/(Γ(*k*)〈*s*〉*^α^*). Note that 〈*P_k_*〉 decays with an exponent less than *k* – 1; therefore the mean probability of parallel evolution is asymptotically much larger than in the case of light-tailed DBFEs. The scaling *n*^-(*α*–1)^ in (4) was first reported in [9] and recently derived independently in [13] in a different context. In particular, we see that when 1 < *α* < 2, (*P*_2_) decays anomalously, *i.e*. with an exponent < 1, in contrast to *P*_2_ ~ *n*^-1^ as in the light-tailed case; see Fig 1(a).

It is important to point out that *P_k_* does not become sharply centred around its mean value when *k* > *α*, which can be shown as follows. The *m*-th moment is given by 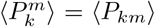 (see SI). The value of 〈*P_km_*〉 can be read off from (4) by replacing *k* by *km*. Thus, all moments are of the same order *n*^-(*α*-1)^. In particular, we notice that for 1 < *α* < 2, 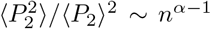. For self-averaging systems (which obey (3) for all *k*), this ratio goes asymptotically to 1, and the standard deviation vanishes relative to the mean. In contrast, here we see that the standard deviation diverges relative to the mean, implying a broad distribution for *P*_2_, as illustrated in Fig 1(b). This non-self averaging effect arises because the sum 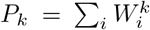 is dominated by the largest weight 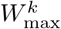 [13, 14]. According to EVT, the highest selection coefficient scales as *n*^1/*α*^, implying that *W_max_* ~ *n*^1/*α*-1^. Therefore, the scale of typical *P_k_* is

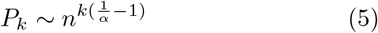

for *k* > *α*, which is asymptotically smaller than (*P_k_*) as given by (4). In fact, most of the weight is concentrated near the typical value, and the much higher mean is obtained from values of *P_k_* that are much rarer but have much higher magnitude.

### Case II

For the severely heavy-tailed case 0 < *α* < 1, all integer moments of s diverge. It was shown in [14] that for a power-law distribution with 0 < *α* < 1,

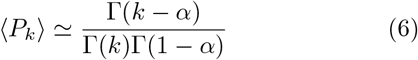

in the limit of large *n*. Specifically, the average probability of parallel evolution in two replicates is 〈*P*_2_〉 ≃ 1 – *α*. Note that the asymptotic form in (6) is independent of *n*, and thus we have the striking result that the probability of parallel evolution remains finite even in the limit of an infinite number of available alternative mutations. In the present case, all moments of *P*_2_ are of *O*(1), and therefore *P*_2_ is non-self averaging. This is visible in the wide distribution of *P*_2_ as shown in the numerically sampled plot in the inset of Fig 1(b). Similar non-self-averaging effects are familiar in the physics of disordered systems (see [14] and references therein), and in probability theory [15].

While the moderately (*α* > 1) and severely (*α* < 1) heavy-tailed cases display somewhat different behavior, we note that both (4) and (6) give rise to the recursion relation

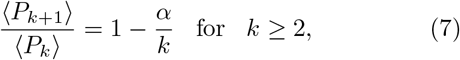

which therefore holds for the entire range 0 < *α* < *k*. The result is independent of *n* and of all features of the underlying distribution except the tail exponent *α*. It is therefore suitable for extracting *α* from empirical data; however, the disadvantage is that the averages require large datasets. Equation 7) easily yields an approximate solution for large *k*, 〈*P_k_*〉 ~ 1/*k^α^*, which is verified in Fig 1(c). The slow decay of 〈*P_k_*〉 with *k* contrasts with the exponential decay of the *typical P_k_* as given by (5).

To summarize the theory, we firstly note that for a heavy-tailed DBFE, the probability of parallel evolution is asymptotically higher than that for light-tailed distributions. The second important observation is that *P*_2_ varies widely even across large, independent samples generated from the same heavy-tailed DBFE. This counter-intuitive property arises because *P*_2_ is dominated by the weights *W_i_* of only a few mutations even as *n* → ∞, which also implies that knowledge of the DBFE does not allow us to predict the degree of repeatability of evolution. Rather, one needs to know the values of the few dominant selection coefficients for the system in question.

## III. APPLICATIONS

Empirical estimates of the tails of the DBFE are available for a number of microbial strains in different environments. Several studies have found evidence of distributions with finite moments [8]. The precise distribution is often difficult to obtain, and some authors prefer to follow the extreme-value hypothesis and estimate the EVT class to which the DBFE belongs [7, 9, 10]. Several studies have uncovered evidence for heavy-tailed DBFEs. For example, studies with viruses [11] and yeast [10] found the fitness effects to be heavy-tailed in particular environments, with *α* > 2.

The theoretical results discussed in this article are valid asymptotically, *i.e* in the limit of large n. This makes the application of the theory to empirical data a challenging task. Nonetheless, we will show that signatures of non-self-averaging effects can be discovered in empirical data sets. For this purpose, we use data on selection coefficients associated with antibiotic resistance evolution reported in [9]. In this study, the fitness effects of 48 beneficial mutations in the resistance enzyme TEM-1 *β*-lactamase were reported for *Escherichia coli* growing at four different concentrations of the antibiotic cefotaxime (see SI for further details of the experiment). An analysis based on EVT indicated that the DBFE belonged to the Gumbel class (light-tailed) for the lowest concentration [9]. At the three higher concentration, the analysis indicated power-law distributions, though large uncertainties were associated with the inferred values of the exponent [9].

We evaluate the statistics of *P*_2_ for the four different concentrations (see SI for further details). Fig 1(d) shows *P*_2_ as a function of n. For the lowest concentration, *P*_2_ is seen to be small with a small dispersion, and it decreases with *n*, consistent with our expectation. For the three higher concentrations, the values of *P*_2_ are larger and have a large dispersion, which is consistent with heavy tailed distributions. There is no discernible decrease with n. However, due to the relatively small values of *n* and the modest size of the data sets, it is not possible to distinguish this from a slow decrease with n.

In Fig 1(e), we have plotted *P_k_* as a function of *k*. Note that the distinction between the typical and mean values [13] has important implications here. Due to the limited size of the data, we have not used the recursion relation (7) to infer α. Instead, for each concentration, we have used the entire set of selection coefficients to create a single sample value of *P_k_*, which is expected to be of the typical scale given by (5). Using this, we estimate the exponent α which is seen to progressively decrease with increasing concentration. indicating an increasingly heavy-tailed distribution. Thus, stronger selection pressures amplify the differences between fitness effects of beneficial mutations, leading to a broader distribution. Nonetheless, we should mention that inferred power-law exponents should be treated with some caution, since these can be sensitive to experimental errors or methods of analysis [9, 11]. What is clear from Fig 1, however, is that the behavior of *P_k_* is at least a good qualitative indicator of the dispersion of selection coefficients. We have also computed and plotted the effective mutation number *n_e_* in Fig 1(f). The trend is again seen to be as predicted by theory. At the lowest concentration, n_e_ is relatively large and close to (*n* + 1)/2 (where *n* is the actual number of mutations), consistent with an exponential distribution of selection coefficients. The effective mutation number decreases progressively with increasing concentration, and indicates a slower than exponential tail.

## Acknowledgements

SGD and JK acknowledge support by the Deutsche Forschungsgemeinschaft (DFG, German Research Foundation) within SFB 1310 *Predictability in evolution*.

## Supplementary Information

### Probability of fixation of beneficial mutations

The probability of fixation of the *i*-th mutation is proportional to *s_i_* when *s_i_* is positive but small. Denoting *F_i_* as the fitness of the *i* – th mutant arising in the background genotype with fitness *F*_0_, we have 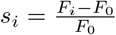 (by definition). We require this quantity to be small, even though the fitness effects *S_i_* ≡ *F_i_* – *F*_0_ are chosen from a distribution supported over the half-line [0, ∞), which means that *S_i_* can be arbitrarily high. Thus, for the small s approximation to hold, *F*_0_ must be large enough such that, with high probability, the largest of *n* chosen values of s is much smaller than *F*_0_. Let the distribution of the largest value be *P_e_*(*s; n*), and we define 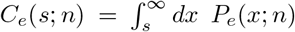. We therefore require that *C_e_*(*ϵF*_0_; *n*) ≪ 1, where *ϵ* ≪ 1. For power law tails of *P*_s_(*s*) as considered here, it is known that 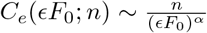, and therefore the approximation holds when 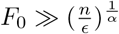.

### Tails of empirical DBFEs

Empirical estimates of the tails of DBFEs are available for a number of different systems. Below we discuss reports in the literature of both light and heavy-tailed DBFEs.

#### Light-tailed DBFEs

Several studies have found evidence of distributions with finite moments. For example, evidence of exponential distributions were found for *Pseudomonas fluorescens* [16] across a range of environments, and for *Pseudomonas aeruginosa* [17] at low antibiotic concentration. A normal distribution was reported [18] for *P. fluorescens* in an environment where the ancestral type has extremely low fitness. While the normal distribution is light-tailed, it is not a limiting EVT distribution (which is not particularly surprising here [18], since the EVT hypothesis assumes that the ancestral type is relatively well-adapted). The DBFE for two bacteriophage viruses have been found [19] to be consistent with the uniform distribution, which is a special case of the limiting Weibull distribution of EVT. The DBFE for a protein in the hepatitis C virus was found [20] to be consistent with an exponential distribution. A study [10] on the protein Hsp90 in the yeast *Saccharomyces cerevisiae* determined the DBFE across several environments and estimated it to be of the Weibull type in all but one environment.

#### Heavy-tailed DBFEs

A number of experiments have uncovered evidence for heavy-tailed distributions as well. For example, a study with an antibiotic resistance enzyme in *E. coli* [9] found evidence of heavy-tailed distributions at high antibiotic concentrations, with *α* belonging to both the moderately (*α* > 1) and severely (*α* < 1) heavy-tailed regimes. At low antibiotic concentration, the same study estimated the DBFE to be light-tailed. This is consistent with the previously mentioned work with *P. aeruginosa* [17] which reported that, at high drug concentration (and therefore low wild type fitness), the DBFE was broader and could not be fit with an exponential distribution. A study [11] on the influenza A H1N1 virus inferred that the DBFE belongs to the Weibull domain in the absence of the antiviral drug oseltamivir and to the Fréchet domain in its presence. The previously discussed work on Hsp90 in yeast [10] found the fitness effects to be heavy-tailed, with *α* > 2, in an environment with lowered temperature and elevated salinity. in a related study [21], the DBFE of synonymous mutations in Hsp90 was found to be heavy-tailed in several environments with *α* > 2 in most cases. A heavy-tailed DBFE has also been detected among mutations in tumors [12]. A common pattern in many of these results is that the DBFE becomes broader in novel and challenging environments.

### Number of available beneficial mutations

The fraction of all mutations (occurring before the biasing effect of selection) that are beneficial is expected to vary strongly across organisms and environments, and is difficult to determine. Nevertheless, studies based on laboratory experiments with microbes have often concluded that beneficial mutations are relatively common. For example, Schenk et al. [9] estimate that 3.4 % of all base pair substitutions in the TEM-1 *β*-lactamase gene increase resistance to cefotaxime, and two survey articles [22, 23] state that the typical beneficial fraction among genome-wide mutations in bacteria is on the order of 10^-2^. The *E. coli* genome has about 4.6 × 10^6^ base pairs [24]. Therefore, even if one considers only point mutations, there are potentially about 10^4^ beneficial mutations.

### Probability of parallel evolution for exponential distributions

Following [14], we notice that, by definition,

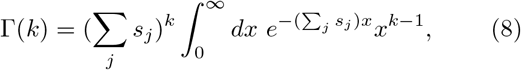

and therefore we have

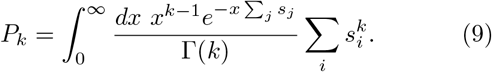

The mean probability is

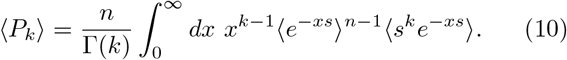

Note that one can simply use the exponential distribution *P_s_*(*s*) = *e^-s^*, since a rescaling of the distribution does not alter the homogeneous expression for *P_k_*. It is easy to show that for this distribution, 〈*e^-xs^*〉 = 1/(1 + *x*) and 〈*s^k^e^-xs^*〉 = Γ(*k* + 1)/(1 + *x*)^*k*+1^. Substituting these in equation (10) we get

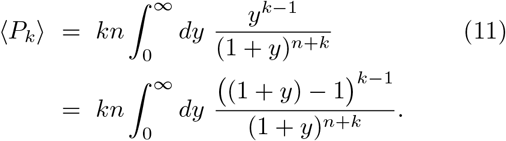

Performing the binomial expansion of ((1 + *y*) – 1)*^k^* and evaluating the subsequent integrals leads to

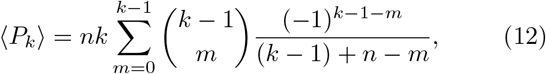

which reduces to the result of Orr [6] for *k* = 2.

### Mean probability of parallel evolution for moderately heavy-tailed distributions

Here we obtain the asymptotic expression of 〈*P_k_*〉 for *k* > *α* > 1. A closely related proof is available in [13]. For large *n*, the integral in (10) is dominated by the small x behavior, and for sufficiently small *x* we have

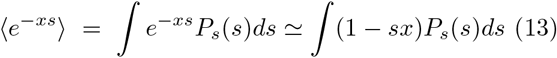

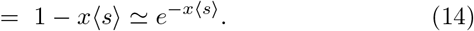

Further, for *k* > *α* [14],

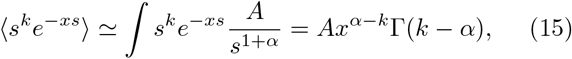

where the first step is justified because, for sufficiently small *x*, the dominant contribution to the integral comes from the tail of the distribution. We now use (14) and (15) in (10) to obtain the result in the main text:

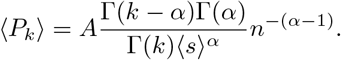

### Moments of the probability of parallel evolution for moderately heavy-tailed distributions

The *m*-th moment of *P_k_* is given by

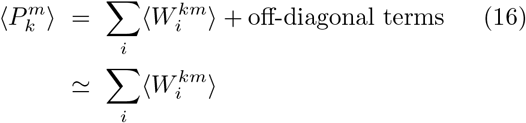

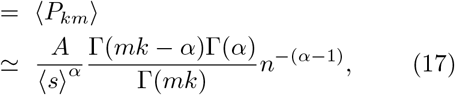

where the second term in (16) is dropped because, as we show now, it is of sub-leading order. The off-diagonal terms in 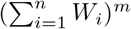 can be written as 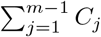 where

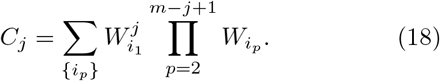

The sum has 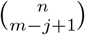 terms, which, in the limit of large *n*, is *O*(*n*^*m-j*+1^) terms. Using the form (8) of the Γ-function, we write

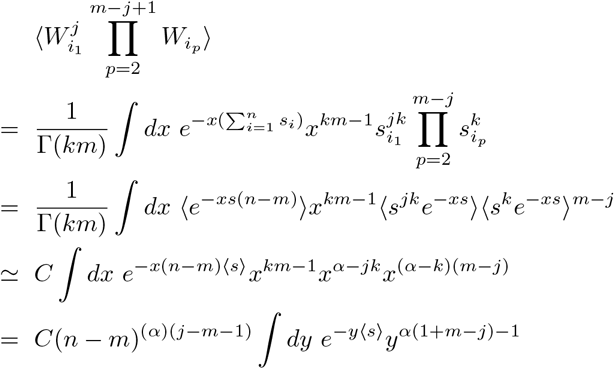

where *C* = *A*^1+*m-j*^Γ(*jk* – *α*)(Γ(*k* – *α*))^*m-j*^/Γ(*km*). Note that the *n*-dependence is *n*^-*α*(*m-j*+1)^, and since there are *O*(*n*^*m-j*+1^) such terms, *C*(*j*) ~ *n*^-(*α*-1)(1+*m-j*)^. Since *j* ≤ *m* – 1, the exponent (*α* – 1)(1 + *m* – *j*) > *α* – 1, and therefore *C_j_* decays faster than the leading term given in (17).

### Numerical simulations

For Figs 1(a)-(c) of the main text, large random samples were generated from heavy-tailed distributions. We used a Monte Carlo procedure outlined in [14]. We recapitulate it here for completeness. First, we generate independent random numbers {*x*_1_, *x*_2_,… } distributed uniformly between 0 and 1. Then we write 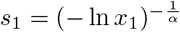, and use the recursion relation

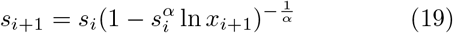

to generate the sequence {*s*_1_, *s*_2_,… }. The value of *P_k_* calculated from this sequence has the desired statistics.

### Details of Data Analysis

The analyzed data was obtained from the study reported in [9].

### Brief description of the relevant experiments in Ref. [9]

In this study, PCR mutagenesis was used to introduce random mutations into TEM-1 *β*-lactamase on plasmids in *Escherichia coli*. 48 mutations with beneficial effects on cefotaxime (CTX) resistance were identified. Fitness at various cefotaxime concentrations were inferred from survival data using a branching model. Selection coefficients were computed relative to the least fit mutant at each CTX concentration.

### Data Analysis

We have used the selection coefficients from [9] in our study. The expression for *P_k_* used for analysis was

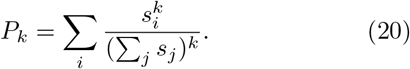

In Fig 1(d), we split the total dataset of the selection coefficients randomly into disjoint subsets at each concentration. Each subset was used to produce one value of *P*_2_, and the boxplots were created from the values of *P*_2_ obtained in this way. For Fig 1(e), the entire dataset at each concentration was used to obtain the value of *P_k_*. Error estimates were not available for the selection coefficients from [9]. However, measurements of resistance level to cefotaxime in [9] had a mean error level of approximately 10 per cent which was used as a rough estimate of the error level in the selection coefficients. The dominant contribution to the variation in *P_k_* comes from the numerator of (20) and this was taken to be the only source of error as an approximation. Subsequently, standard error propagation was used to estimate the error in *P_k_* from that in the selection coefficients. The values of *α* were obtained by non-linear least squares fitting of the data. In Fig 1(f), the actual mutation number was reported as the number of selection coefficients used for our analysis. The error bars in the effective mutation number were obtained through error propagation from *P*_2_.

